# The antioxidant drug N-acetylcysteine abolishes SOS-mediated mutagenesis produced by fluoroquinolones in bacteria

**DOI:** 10.1101/428961

**Authors:** Ana I. Rodríguez-Rosado, Estela Ynés Valencia, Alexandro Rodríguez-Rojas, Coloma Costas, Rodrigo S. Galhardo, Jesús Blázquez, Jerónimo Rodríguez-Beltrán

**Affiliations:** Instituto de Biomedicina de Sevilla (IBiS), Seville, Spain.; Department of Microbiology, Institute of Biomedical Sciences, University of São Paulo, São Paulo, Brazil.; Institute of Biology, Freie Universität Berlin, Berlin, Germany.; Centro Nacional de Biotecnología (CNB), Madrid, Spain

## Abstract

Certain antibiotics, particularly fluoroquinolones, induce the mutagenic SOS response and increase the levels of intracellular reactive oxygen species (ROS), which have been associated with antibiotic lethality. Both SOS and ROS promote bacterial mutagenesis, fueling the emergence of resistant mutants during antibiotic treatments. However, the relative contribution of ROS and SOS on this antibioticmediated mutagenesis is currently unknown. We used the antioxidant molecule N-acetylcysteine (NAC) to study the contribution of ROS on the SOS response and the mutagenesis mediated by the fluoroquinolone anti-biotic ciprofloxacin (CIP). We show that NAC is able to reduce intracellular ROS levels, as well as the SOS response caused by treatment with subinhibitory concentrations of CIP, without affecting its anti-bacterial activity. This effect reduces anti-bioticinduced mutagenesis to levels comparable to a translesion synthesis DNA-polymerases deficient strain, suggesting that ROS play a major role in SOS-induced mutagenesis. Collectively, our results shed light on the mechanisms underlying antibioticinduced mutagenesis and open the possibility for the use of NAC as adjuvant in antibiotic therapy to hinder the development of antibiotic resistance.

## Importance

The development of antimicrobial resistance, together with the existing paucity in the antibiotic pipeline, renders every antibiotic into a non-renewable resource that should be carefully rationed. This worrisome scenario is exacerbated by the fact that treatment with certain antibiotics, besides killing bacteria, increase the chances of surviving bacteria to acquire resistance as a side-effect. The mechanisms underlying this phenomenon involve complex bacterial physiological responses to antibiotics such as induction of the SOS response and the generation of reactive oxygen species. In this work, we demonstrate that the antioxidant drug N-acetylcysteine inhibits antibiotic-induced mutagenesis by reducing the levels of reactive oxygen species and SOS induction in bacterial cells upon antibiotic treatment. Our results strongly suggest that reactive oxygen species are a key factor in antibiotic-induced SOS mutagenesis and open the possibility of using NAC combined with antibiotic therapy to counteract the development of antibiotic resistance.

## Introduction

Antibiotics, besides their antimicrobial action, can promote genetic variability in bacteria as an undesirable side effect (1). In turn, genetic variability increases the chances for bacteria to acquire resistance and jeopardize the success of antimicrobial therapies. Blocking the bacterial physiological responses that promote genetic variability is thus crucial to hinder resistance spread (2, 3).

Most of the genetic variability produced by antibiotics has been attributed to the induction of the SOS response. The SOS response is a coordinated genetic network that responds to DNA damage. Several antibiotics produce DNA damage and consequently induce the SOS response. For instance, fluoroquinolones such as ciprofloxacin (CIP) block DNA gyrase on DNA, which cause the stalling of replication forks and leads to cell death (4). This produces double strand breaks (DSB) that are processed into single strand DNA, that together with RecA trigger the SOS response. SOS induction up-regulates the expression of more than 40 genes whose functions include DNA-damage tolerance and non-mutagenic DNA repair (5–8). However, when DNA damage is persistent, the error-prone DNA translesion synthesis (TLS) takes place. In *Escherichia coli*, TLS is accomplished by the specialized DNA polymerases Pol II, Pol IV and Pol V, encoded respectively by the *polB*, *dinB* and *umuDC* genes (9). TLS polymerases are able to replicate heavily damaged DNA but do so at the cost of a reduced fidelity, therefore increasing mutagenesis (5). Additionally, RecA-mediated recombination is also induced by fluoroquinolone antibiotics (10). Hence, some antibiotics can promote mutagenesis and recombination (i.e. genetic instability) by directly inducing DNA damage and, in turn, the SOS response.

Bactericidal antibiotics have been shown to produce a perturbation of the intracellular redox homeostasis. This perturbation is caused by an increased intracellular respiration rate accompanied by destabilization of the Iron-Sulfur clusters, which leads to production of reactive oxygen species (ROS) via Fenton chemistry (11–13). ROS are highly reactive chemical species capable of rapidly oxidizing key cellular components, including proteins, lipids and DNA. ROS have been argued as a common cause of bacterial cell death for several antibiotic families (12–14), although this notion has been further challenged (15, 16). Oxidation of DNA by ROS produces a wide variety of lesions that, if not repaired, are mutagenic and can even cause cell death (17–19).

In summary, previous studies have shown that there are at least two routes to antibiotic-triggered bacterial mutagenesis; SOS-mediated TLS and ROS-induced mutagenesis. Interestingly, these two routes are probably not independent but highly intertwined. For instance, oxidation of the nucleotide pool after antibiotic treatment leads to Pol IV-mediated incorporation of 8-oxo-dGTP into DNA, which creates a mutagenic lesion (14, 20). Furthermore, ROS are good SOS inducers because they directly damage DNA (21–23). Hence, SOS-mediated TLS mutagenesis might be fueled by the presence of oxidative damage in both DNA and the nucleotide pool (24, 25).

Recently, there has been growing interest in the development of novel therapies designed not to kill bacteria, but to inhibit the aforementioned routes to antibiotic resistance (2, 26, 27). In particular, most studies have focused on developing SOS inhibitors that target RecA (28–30). RecA offers an appealing target because its inhibition not only reduces bacterial evolvability, but also renders bacteria more sensitive to several antibiotics (31); in some cases leading to complete reversion of antibiotic resistance (32). However, the presence of RecA homologs in mammals in humans suggests LexA as a potentially safer target (33).

In this study, we hypothesized that antioxidant molecules could reduce ROS produced by antibiotic treatment and consequently inhibit SOS induction. This combined inhibition might, in turn, reduce antibiotic-induced mutagenesis. To test this idea, we focused on N-acetylcysteine (NAC), a well-known antioxidant that acts as a scavenger of oxidant species and as a precursor of glutathione synthesis (34, 35). NAC is clinically safe and is currently used in humans therapy to treat numerous disorders (34). Additionally, NAC does not negatively affect the activity of major antibiotic classes, with the exception of carbapenems (36, 37). On the contrary, NAC has shown antimicrobial properties against a range of clinically-relevant pathogens (38–41). Our results show that, due to its antioxidant properties, NAC offers an unique opportunity to disentangle the effects of ROS and SOS in antibiotic-mediated mutagenesis. Additionally, it promises to become a therapeutic alternative to outsmart the evolution of bacterial resistance.

## Results

### CIP induces mutagenesis at subinhibitory levels

The mutagenic activity of antimicrobials is expected to occur within a window of concentrations around their minimal inhibitory concentration (MIC), because higher levels would kill cells or stop their growth while lower concentrations would not have a stimulatory effect (42). To determine the concentration of CIP that induces the highest increase in mutagenesis, we treated exponentially growing cultures of *E. coli* strain IBDS1 (43) with different concentrations of CIP ranging from 0.25 to 4 times the MIC for 8 hours. We then determined mutation rates using two independent selective markers. Cells treated with 8 ng/ml of CIP (which correspond to ½ of the MIC) showed the highest increase on the rate of mutations conferring resistance to rifampicin (Rif-R; 24-fold increase) and tetracycline (Tet-R; 2.5-fold increase) (**Supplementary figure 1**). Higher concentrations of CIP barely produced any increase in mutagenesis. This probably occurred because these concentrations hampered growth of most treated cells (**Supplementary figure 2**), drastically reducing effective population size and hence limiting evolvability (44). We decided to use 8 ng/ml of CIP hereafter to maximize antibiotic-induced mutagenesis.

### NAC reduces ciprofloxacin-induced intracellular ROS

We then measured the levels of ROS caused by treatment with CIP and tested if NAC was able to reduce CIP-generated ROS. To this end we used 2′,7′-Dichlorofluoresce indiacetate (H_2_DCFDA), a ROS sensitive dye that emits fluorescence when it is oxidized intracellularly (45). H_2_DCFDA has been previously shown to be an extremely sensitive probe for the detection of ROS caused by fluoroquinolones, detecting with great sensitivity H_2_O_2_, ROO· and ONOO^-^ (45). As expected, CIP consistently showed increased fluorescence levels over those produced by antibiotic-mediated autofluorescence (46), indicating a massive increase of ROS levels compared to untreated cells (**Figure 1** and **Supplementary figure 3a**). Most important, the induction of ROS was reverted to nearly basal levels when CIP treatment was combined with NAC at 0.5%. This result suggests that NAC, at a physiologically attainable concentration (47), might be able to reduce DNA damage by reducing the levels of ROS upon CIP exposure.

### NAC reduces CIP-mediated induction of the SOS response

We then analyzed the effect of CIP and NAC in SOS induction. To this end, we used the strain IBDS1 pRecA::*gfp* that harbours the transcriptional fusion P*recA*::*gfp* contained in a low copy number plasmid (48, 49). As expected, CIP strongly induced the SOS response approximately 14-fold compared to untreated controls (Tukey multiple comparisons after significant ANOVA, P>0.001; **Figure 2a**).

**Figure 1.**
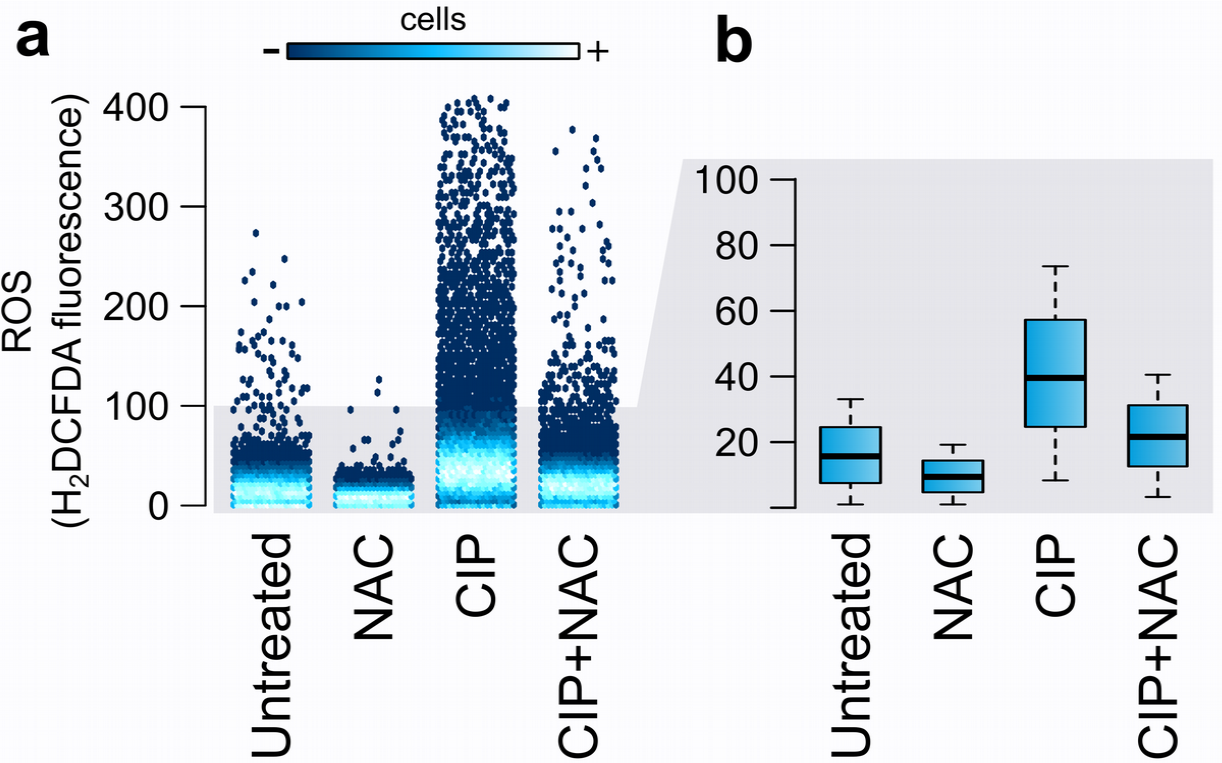
NAC reduces ciprofloxacin-induced intracellular ROS. OS were assessed by individually capturing the fluorescence of H_2_DCFDA in 30,000 cells by flow cytometry after 8 hours of treatment with either 8 ng/ml of CIP, 0.5% NAC or both agents in combination. An untreated control is shown as a reference. a) The dot plot shows the distribution of fluorescence signal in the treated populations. At least 99% of the events recorded are shown. The color scale displays the density of events at every fluorescence level. b) To allow better comparison, the data is depicted as boxplots, in which the horizontal line represents the median value, the depth of the box represents the interquartile range (50% of the population), and the whiskers extend to 0.5 times the interquartile range. Note that shaded areas in both panels represent the same data.

Following the hypothesis that antioxidant compounds can be effective inhibitors of SOS induction (50), we tested the effect of different NAC concentrations on CIP-mediated SOS induction. NAC inhibited SOS induction caused by CIP at all concentrations (Tukey multiple comparisons after significant ANOVA, P>0.001; **Figure 2a**), with an IC50 of 0.5% (**Figure 2c**). Again, this concentration is within the range of attainable physiological values after inhaled administration (47), strongly suggesting that inhibition of the SOS response by NAC could be a feasible therapeutic approach. Importantly, NAC did not reduce CIP bactericidal activity, as stated by MIC results (**Supplementary Table 1**),growth curves (**Figure 2b**)and a checkerboard assay (**Figure 2d**). On the contrary, NAC alone inhibited bacterial growth without inducing the SOS response (Tukey multiple comparisons after significant ANOVA, P=0.0003). This result is in line with previous studies that reported NAC antibacterial properties against a range of bacterial pathogens (37, 40, 41). Additionally, we verified that the differences in the final optical density observed upon different treatments do not directly influence the measurement of SOS induction using GFP fluorescence (**Supplementary figure 4**).

SOS induction leads to the overexpression of *sulA* (*sfiA*), whose product inhibits FtsZ ring formation and hence cell division (51). The phenotypic consequence of cell division inhibition is filamentation, which offers an additional SOS-dependent measurable phenotype. We assessed whether 0.5% NAC was able to inhibit CIP-mediated cell filamentation by both flow cytometry and direct observation of Gram-stained cultures. **Figure 3** shows that, as expected, CIP treatment produces a vast increase in the fraction of the population with filamented cells compared with untreated cultures (33% versus 0.3% of filamented cells). Remarkably, administration of 0.5% NAC together with CIP, prevented filamentation in a large fraction of cells (14% versus 33%, for CIP+NAC versus CIP alone). We qualitatively confirmed these results by microscopy observation of Gram-stained cells (**Figure 3b**).

**Figure 2.**
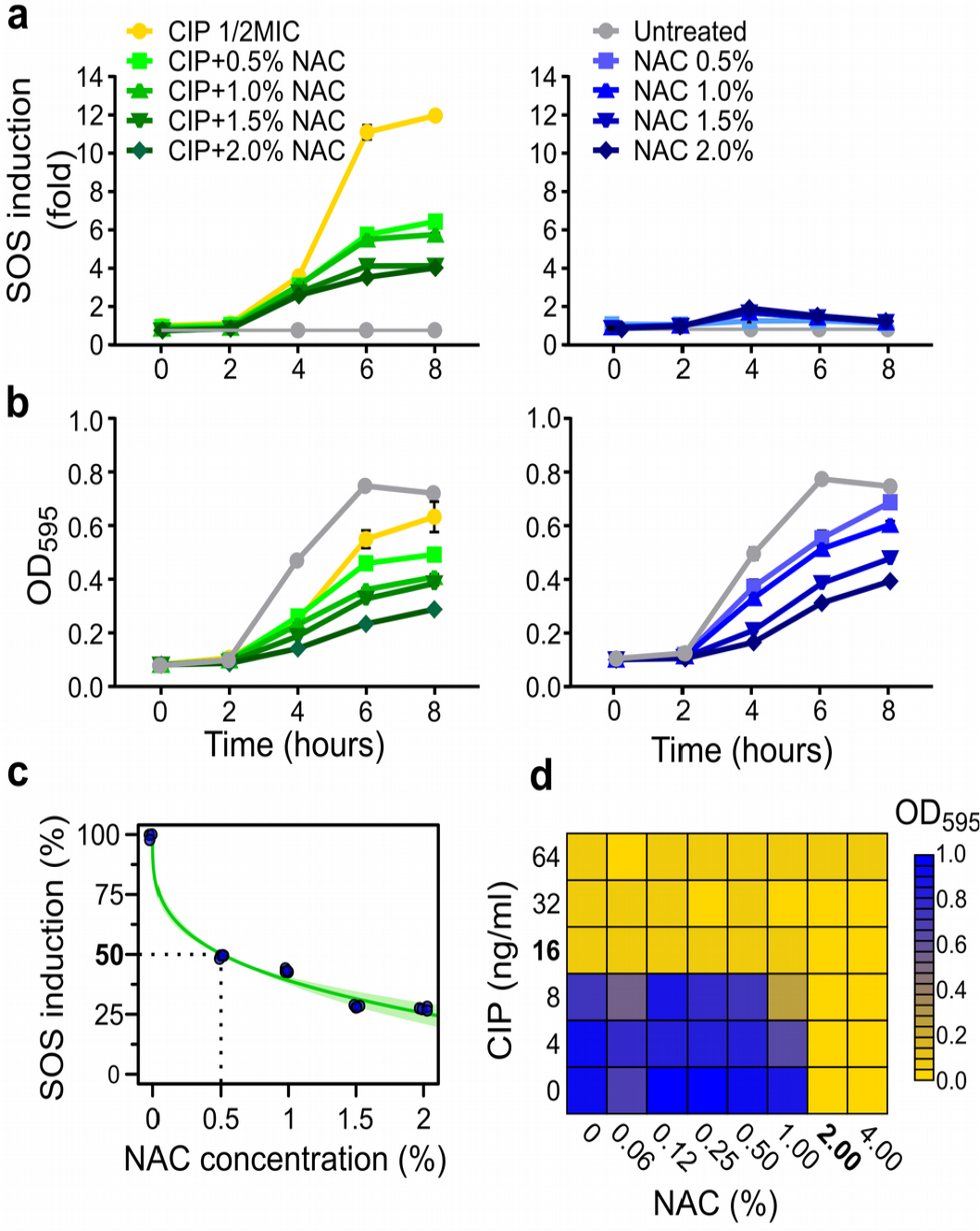
NAC reduces ciprofloxacin-induced SOS response. Bacterial growth and SOS induction were monitored during treatment with varying concentrations of CIP alone or in combination with NAC (left panels), or NAC alone (right panels). Samples were taken at indicated time points and SOS induction (**a**)and absorbance (**b**)were quantified. Error bars represent standard deviation and are not shown when smaller than data points. (**c**)CIP-mediated SOS induction at 8h was assessed in combinations with a range of NAC concentrations giving rise to a dose-response curve. Experimental data was fitted to the following formula SOS=a*NAC^b^ by a non-linear model (nls function in R, R^2^=0.988). The concentration of NAC that inhibits 50% of SOS response (IC_50_) was 0.5%. Green shaded area represents 95% confidence interval of the fit. (**d**)Potential interactions of CIP with NAC were determined by the checkerboard method. MIC concentrations for each compound alone are shown in bold typeface. No synergistic or antagonistic effect was found.

In summary, these results demonstrate that NAC does not decrease bacterial susceptibility to CIP. It does, however, significantly reduce up to a 75% CIP-mediated induction of *recA* transcription and cell filamentation, hallmarks of SOS induction.

**Figure 3.**
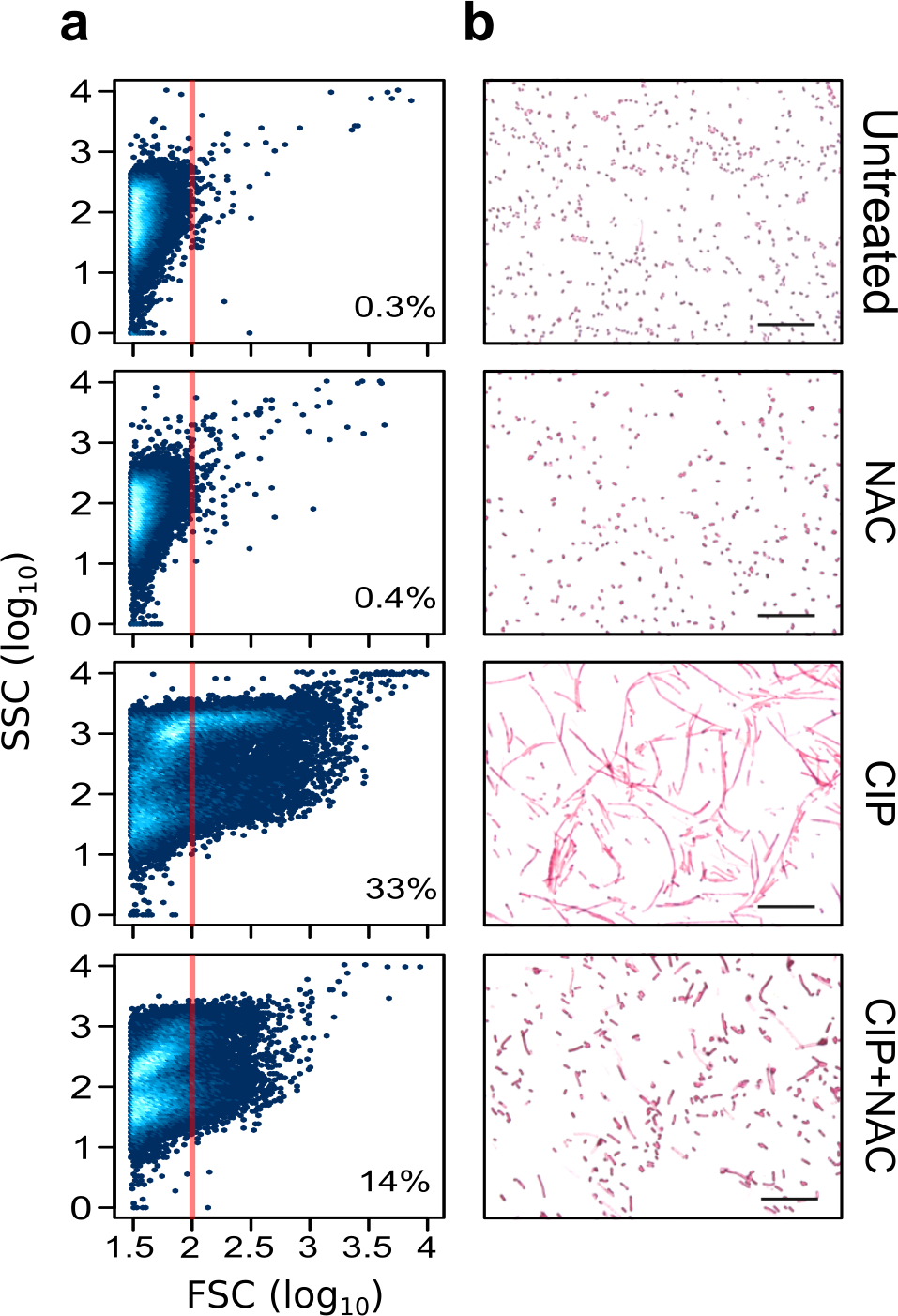
NAC reduces CIP-induced filamentation. **a**)The fraction of filamented cells after the stated treatments is shown by means of flow cytometric analysis of 30,000 cells (SSC; side scatter FSC; forward scatter; proportional to cell size). The percentage on every graph represents the filamented fraction of the population (log_10_(FSC)>2; red vertical line). (**b**)Representative microscopy fields of Gram-stained cells. Scale bars is 20*μ*m.

### NAC inhibits SOS response in a ROS-dependent manner

Although the above results compellingly suggest that the reduction of CIP-induced ROS underlie SOS inhibition by NAC, we cannot rule out other possibilities. NAC could potentially perturb the activity of the SOS regulatory machinery, for example inhibiting RecA-ssDNA nucleation or LexA self-cleavage. If that were the case, we expect that NAC would reduce SOS induction also when DNA damage is independent of ROS. To test this possibility, we used the *E. coli* strain SMR14354 (52), whose chromosome carries a unique cutting site for the restriction enzyme I-SceI. In the presence of 0.1% L-arabinose (Ara), I-SceI is produced, generating DSB and consequently inducing SOS response (**Figure 4a**). Using flow cytometry and H_2_DCFDA we first verified that generation of DSB by I-SceI does not increase ROS levels as a side effect (**Figure 4b** and **Supplementary figure 3c**). We then measured SOS induction at different time-points after DSB induction. Our results demonstrate that the addition of NAC causes no measurable inhibition of the SOS response (Two tailed Student’s *t* test, t=0.64, df=4, P=0.56 for Ara vs Ara+NAC after eight hours of treatment), indicating that NAC inhibition of the SOS response is ROS-dependent (**Figure 4c**). Although the experimental conditions used here have been shown to cause at least a single DSB in 90% of the cells (and more than one in 50% of cells) (52), induction of the SOS response is lower than that caused by 8 ng/ml of CIP (~6 versus ~14 fold). An alternative explanation for our results could be that at lower SOS inductions, NAC is unable to decrease SOS induction. To discard this possibility, and to match the level of induction caused by I-SceI mediated-DSB, we tested the effect of NAC in cultures treated with lower CIP concentrations. Our results show that NAC is able to reduce CIP-induced SOS response in all cases (**Supplementary figure 5**).

**Figure 4.**
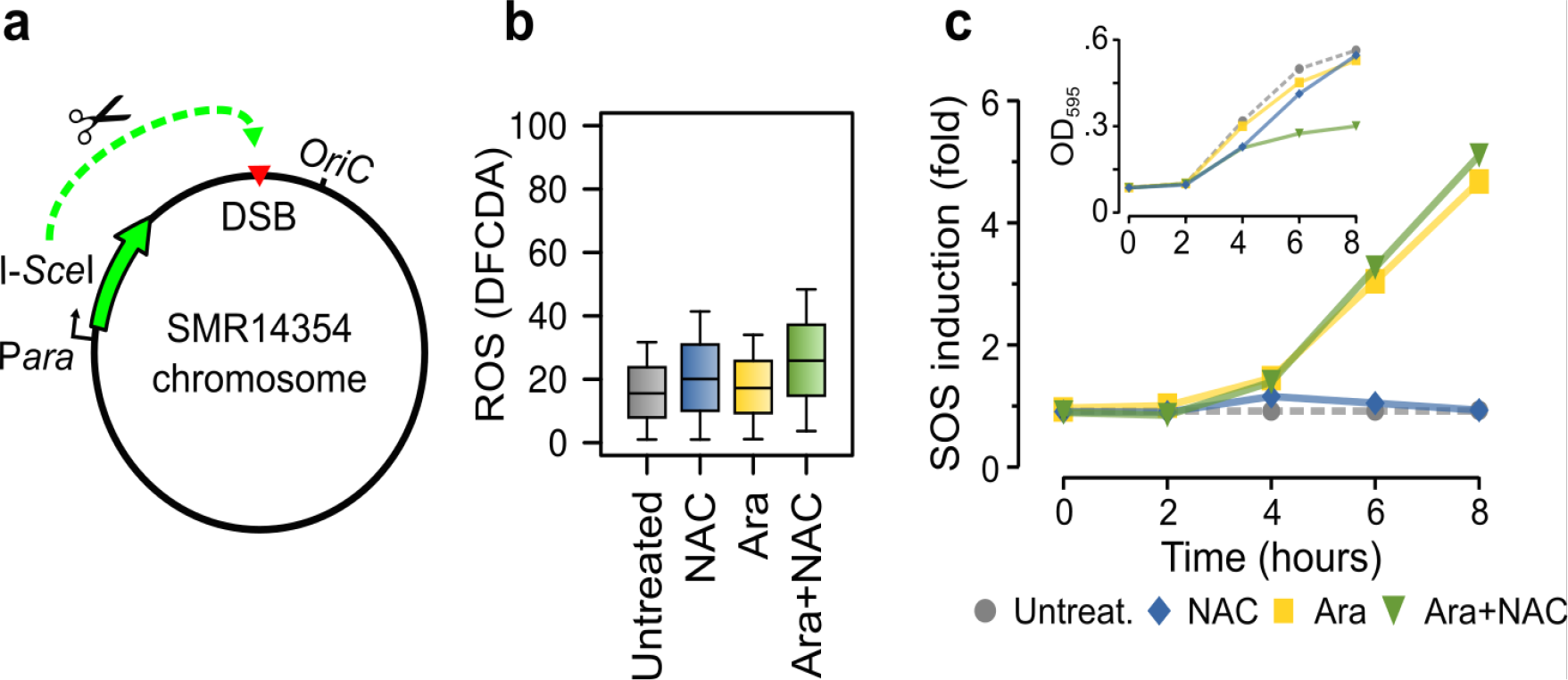
Artificially generated double-strand breaks induce the SOS response but do not generate ROS. a) Schematic diagram of the experimental setting. To generate in vivo DSB, the strain SMR14354 carries a unique cutting site (red triangle) close to OriC of the restriction enzyme I-SceI (scissor), whose expression is induced by L-arabinose (Ara). b) The addition of 0.1% L-arabinose generates DSB that concomitantly induce the activation of the SOS response (yellow curves), measured here by means of an PrecA::gfp transcriptional fusion. The addition of NAC alone (blue) or in combination with Ara (yellow) does not alter SOS induction. Error bars (sd) smaller than data points are not shown for the sake of clarity. Inset graph represents optical density (OD_595_) under the same conditions. c) ROS levels detected by the use of the fluorescent H_2_DCFDA probe and flow cytometry show no significant increase after the induction of DSB for eight hours.

### NAC reduces the SOS-mediated mutagenesis promoted by CIP

The quinolone-mediated increase in mutagenesis has been attributed to the activity of TLS DNA-polymerases, whose transcription is induced as part of the SOS response (9, 53). However, our results and previous studies strongly suggest that high levels of ROS are also mutagenic (17–19). To gain knowledge on the contribution of each of these two mechanisms we used the strain IBDS1 and its TLS^-^ derivative which lacks the three TLS error-prone DNA-polymerases (43). We verified that the TLS^-^ strain showed similar SOS induction and ROS production levels to the wild-type strain when treated by CIP and NAC alone or in combination (**Supplementary figures 3 and 6**). Mutation rates of treated cultures showed that treatment with CIP induced mutagenesis in both WT and TLS^-^ strain, although at lower levels in the TLS^-^ strain (**Figure 5**). This result indicates that a fraction of CIP-mediated mutagenesis is not dependent on SOS TLS-polymerases. The combined treatment with CIP and NAC decreased up to 40% CIP-mediated mutagenesis in the wild-type strain for both Rif-R and Tet-R selective markers (**Fig. 5a**). This highlights the importance of ROS as a major contributor to CIP-induced mutagenesis. On the contrary, NAC was unable to alter CIP-induced mutagenesis in the TLS^-^ strain, indicating that TLS-independent mutagenesis is also ROS-independent (**Fig. 5b**).

Together, these results suggest that treatment in wild-type *E. coli* TLS polymerases act synergistically with ROS in a highly intertwined mutagenesis pathway. Accordingly, reduction of ROS by NAC completely abolishes SOS-mediated mutagenesis in CIP-treated bacteria.

**Figure 5.**
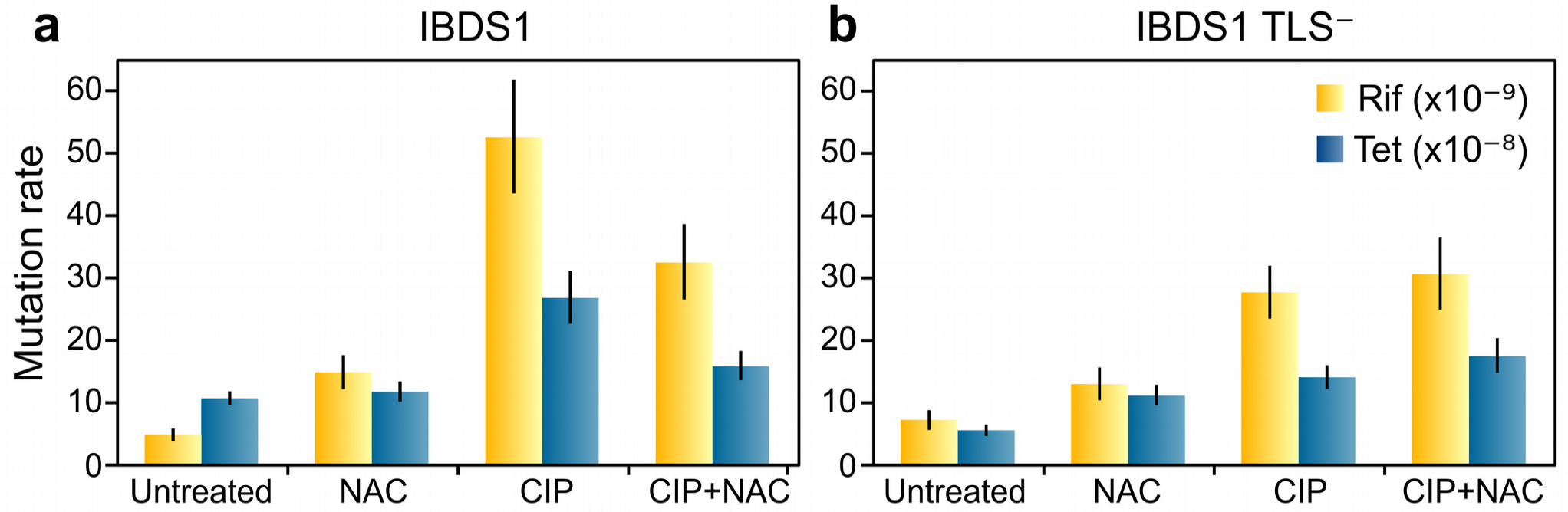
NAC reduces CIP-induced mutagenesis in the wild-type but not in its TLS^-^ derivative. Wild-type (a) and TLS^-^ cells (b) were treated with 8 ng/ml of CIP and 0.5% of NAC alone or in combination. After 20 hours of recovery in antibiotic-free medium, mutation rates (mutations per site per generation) were calculated using rifampicin (yellow bars) or tetracycline (blue bars) as selective markers. Differences are statistically significant when error bars (95% CI) do not overlap.

## Discussion

Fluoroquinolones such as CIP provoke the blockage of DNA gyrase on DNA causing the stalling of replication forks, which produces DSB and leads to cell death (54). DSB are processed to single strand DNA which activates the SOS response causing the upregulation of the SOS genes, including the error-prone TLS DNA polymerases (8, 9). Because TLS DNA replication is mutagenic (9, 53), fluoroquinolones promote mutagenesis by directly inducing DNA damage and, as a result, the SOS-controlled TLS. Additionally, bactericidal antibiotics such as fluoroquinolones increase the intracellular levels of ROS, causing cell death (12, 13). Moreover, high levels of ROS, as DNA-damaging agents, can additionally induce the SOS response and cause mutagenesis (14, 18, 22).

Here, we showed that the decrease of ROS caused by the treatment with the antioxidant NAC attenuates the induction of the SOS response. However, the magnitude of the effect observed in this study (i.e. a reduction of up to 75% of SOS induction by NAC), suggests that ROS are major contributors to DNA-damage and the subsequent activation of SOS in fluoroquinolone-treated bacteria.

This idea is further supported by our mutagenesis results in which a significant reduction, but not abolition, of CIP-induced mutagenesis was observed upon NAC treatment in the wild-type strain. Most important, NAC treatment reduced wild-type mutagenesis to similar levels to those seen in the TLS^-^ strain, suggesting that the residual CIP-induced mutagenesis is independent of TLS repair. Consistent with this view, we observed an increase in mutagenesis in the TLS^-^ strain when submitted to CIP treatment.

This result agrees with recent work in which an increased frequency of indels was found upon CIP treatment in a TLS^-^ strain (55). Together, these results support the existence of a TLS-independent mutagenic pathway. Pomerantz et al. suggested that Pol I can be highly error-prone at RecA-mediated D-loops produced by DSB repair (56). Hence, it is likely that CIP, by generating DSB, can fuel the creation of these RecA-mediated D-loops and the subsequent Pol I mutagenesis. This mechanism is expected to be ROS-independent and could explain the TLS-independent mutagenesis observed in our study. Further experimentation will be needed in order to contrast this hypothesis.

As a clinical application of our results, we propose that NAC could be used as a promising adjuvant in CIP treatment, and possibly with other quinolones. NAC is a clinically safe, FDA-approved drug widely used for the treatment of numerous disorders (35). We have shown that physiologically attainable concentrations of NAC inhibited SOS induction without compromising CIP bactericidal activity. On the contrary, NAC itself has been shown to present antibacterial properties against *Helicobacter pylori* (40), *Haemophilus influenzae* (37), *Stenotrophomonas maltophilia*, *Burkholderia cepacia* (39) and *Pseudomonas aeruginosa* biofilms (41). From the clinical point of view, inhibition of SOS induction could provide several important benefits besides reducing mutagenesis. For instance, SOS-regulated genes control pathogenic processes such as persistence, tolerance, infection, and expression of toxins or virulence factors (57–59). Additionally, it is well known that bacterial filamentation is a SOS controlled process crucial for the development of some bacterial infections, such as urinary tract infections (60). Therefore, inhibition of filamentation could be a desirable therapeutic target to improve the prognosis of bacterial infections. In this regard, our results also show that NAC significantly reduced the fraction of filamented cells after antibiotic treatment.

Taken together, our results suggest that NAC could be used as adjuvant in antibiotic treatment to inhibit SOS-mutagenesis, reducing the chances for the development of bacterial resistance and decreasing pathogenesis without compromising antibiotic activity.

## Methods

### Bacterial strains, plasmids and media

Mutation rate experiments, as well as growth curves and flow cytometry assays, were performed with the *Escherichia coli* MG1655 *att*λ::cI (Ind -) ΔpR *tet* ∆*ara*::FRT ∆*met*RE::FRT strain (IBDS1) and its derivative deficient in error prone polymerases (TLS-). Mutations that inactivate Δ cI (Ind-) represor gene allow the expression of ΔpR *tetA* gene, which confers resistance to tetracycline (TET) (43). The strain SMR14354 (*E. coli* MG1655 ∆*araBAD*567 ∆*attλ*::PBADI-SceI *zfd*2509.2::PN25*tetR* FRT ∆att*Tn*7::FRT *cat*FRT PN25*tetOgam*-*gfp* I-site was used to measure the SOS induction triggered by DSB. This strain carries a unique I-SceI restriction site close to the chromosomal OriC. I-SceI expression is regulated by the Ara inducible P*ara* promoter (52). The plasmid pSC101-Pr*ecA*::*gfp* (48) was used to monitor SOS induction by fluorescence experiments. Bacterial strains were grown in LB Broth media, supplemented with ciprofloxacin (CIP; at various concentrations), kanamycin (KAN; 30 *μ*g/ml) or 0.5 % V/V of N-Acetylcysteine (NAC) when needed.

Minimal Inhibitory Concentration (MIC), as well as checkerboard assay were performed to determine the interaction between ciprofloxacin and NAC, according to standard susceptibility testing, using LB broth instead of Müller-Hinton. Absorbance at 595 nm was determined using a TECAN Infinite^®^ F200 spectrophotometer after 20h of incubation at 37°C.

### Induction of SOS Response

Three independent overnight cultures of the strain containing the plasmid pSC101-P*recA*::*gfp* were diluted 1:100 in 5 ml of LB supplemented with KAN and grown to exponential phase (OD_600_ 0.5-0.6) at 37°C and 200 r.p.m. Subsequently, the cultures were diluted 1:50 in LB+KAN. 2ml aliquots were treated with various concentrations of CIP (with or without NAC 0.5%) during 8 hours at 37°C and 250 r.p.m. The procedure was repeated with the strain SMR14354, but adding 0.1% V/V of Ara instead of CIP as SOS inducer. We verified that higher Ara concentrations do not increase SOS induction, probably because at 0.1% the concentration of I-SceI is enough to cause DSB in most cells (52) (**Supplementary figure 7**). Controls without treatment were also included in every experiment. Absorbance at 595 nm and green fluorescence (485/520 nm) were monitored using a TECAN Infinite F200 plate reader. SOS induction was obtained by normalizing GFP-fluorescence by the absorbance of each sample. To determine fold change, the average SOS induction of three replicas per condition was divided by the average value of untreated samples.

### Flow cytometry

Intracellular ROS levels were determined by the use of the oxidation sensitive probe 2′,7′-Dichlorofluorescein diacetate (H_2_DCFDA, Sigma-Aldrich). Overnight cultures of the strain IBDS1 or its TLS-derivative were diluted 1:100 in 5 ml of LB media containing H_2_DCFDA 100 *μ*g/ml, and grown to exponential phase (OD_600_ 0.5-0.6) at 37°C and 250 r.p.m. Controls without probe were included to monitor autofluorescence. 2 ml aliquots from 1:50 dilutions from both cultures (with and without H_2_DCFDA) were treated with CIP 8 ng/ml or Ara 0.1%, with or without NAC 0.5%, during 8 hours at 37°C and 250 r.p.m. Three replicas of each condition were included in the assay. Green fluorescence emitted by the intracellular oxidation of the dye was determined using a guava easyCyte cytometer (Millipore). Three replicas of 10,000 events each one, with a concentration of 200-400 cells/*μ*l, were analyzed for each one of the conditions. For estimation of filamentation, forward scatter (FSC) was analyzed in samples without H_2_DCFDA. Data analysis was performed using custom scripts in R (www.R-project.org/).

### Microscopy

Cultures were treated with CIP, NAC or both agents exactly as in the mutation rate experiments. After 8 hours of treatment, a frotis of every sample was prepared as follows: 10*μ*l of culture was spread with a loop on a microscope slide. Samples were fixated by heat, stained with safranin for 1minute and then washed with distilled water. Slides were observed under a Olympus BX61 microscope using the 100× objective.

### Mutation Rate Assays

Three biological replicates of 1:100 dilutions from overnight cultures were grown in LB media to exponential phase (OD_600_ 0.5-0.6) at 37°C and 200 r.p.m. Subsequently, 2ml aliquots from 1:50 dilutions were treated with CIP 8 ng/ml, with or without NAC 0.5% during 8 hours at 37°C and 250 r.p.m. After treatment 1 ml of culture was centrifuged for 6 min at 8,000 r.p.m. Cells were resuspended in fresh LB media and incubated 20 hours at 200 r.p.m to allow resolution of filaments. Appropriate dilutions were plated onto LB-dishes containing tetracycline (TET; 15 *μ*g/ml) or rifampicin (RIF; 100 *μ*g/ml) as selective markers, and LB agar for viable counting. Plates were incubated at 37°C for 24 hours. The expected number of mutations per culture (m) and 95% confidence intervals were calculated using the maximum likelihood estimator, applying the *newton.LD.plating* and *confint.LD.plating* functions that account for differences in plating efficiency implemented in the package rSalvador (61) for R (www.R-project.org/). Mutation rates (mutations per cell per generation) were then calculated by dividing m by the total number of generations, assumed to be roughly equal to the average final number of cells. Differences are considered statistically significant when 95% confidence intervals do not overlap.

### Data Availability

The datasets generated during and/or analysed during the current study are available from the corresponding authors on reasonable request.

## Author contributions

J.B and J.R-B conceived the study. A.I.R, E.Y.V, R.S.G, J.B, A.R-R and J.R-B designed the experiments. A.I.R, E.Y.V and C.C performed the experiments. A.I.R and J.R-B analysed the data. J.B and R.S.G provided reagents and materials. A.I.R, J.B. and J.R-B wrote the manuscript. All authors discussed the results and implications and commented on the manuscript at all stages.

## Acknowledgements

We thank Juan J. Infante and D. Andrade Moreno for critical reading of this manuscript and helpful comments. Strains IBDS1 and SMR14354 are kind gifts of Ivan Matic and Susan M. Rosemberg. J.B. was supported by the Spanish Plan Nacional de I+D+i 2013– 2016 and the Instituto de Salud Carlos III, Subdirección General de Redes y Centros de Investigación Cooperativa, Ministerio de Economía, Industria y Competitividad, Spanish Network for Research in Infectious Diseases; grant REIPI RD16/0016/0009, cofinanced by the European Development Regional Fund “A Way to Achieve Europe” and by Operative Program IntelligentGrowth 2014–2020; and grants FIS PI17/00159 (ISCIII/FEDER,UE) and SAF2015-72793-EXP (AEI/FEDER, UE). R.S.G. was supported by Fundação de Amparo à Pesquisa do Estado de São Paulo, Brazil (FAPESP, grant 2014/15982-6) and Conselho Nacional de Desenvolvimento Científico e Tecnológico, Brazil (CNPq, grant 407259/2013-9). E.Y.V was funded by a postdoctoral fellowship from CNPq (grant 236914/2012-0). J.R.-B. is a recipient of a Juan de la Cierva Fellowship, Ministerio de Economía Industria y Competitividad (FJCI-2016-30019).

